# Molecular pharmacology of selective Na_v_1.6 and dual Na_v_1.6 and Na_v_1.2 channel inhibitors that suppress excitatory neuronal activity ex vivo

**DOI:** 10.1101/2023.08.03.551643

**Authors:** Samuel J. Goodchild, Noah Gregory Shuart, Aaron D. Williams, Wenlei Ye, R. Ryley Parrish, Maegan Soriano, Samrat Thouta, Janette Mezeyova, Matthew Waldbrook, Richard A. Dean, Thilo Focken, Mohammad-Reza Ghovanloo, Peter C. Ruben, Fiona Scott, Charles J. Cohen, James R. Empfield, JP Johnson

## Abstract

Sodium channel inhibitors are used to treat neurological disorders of hyperexcitability. However, all currently available sodium channel targeting anti-seizure medications are non-selective among the Na_V_ isoforms which potentially limits efficacy and therapeutic safety margins. XPC-7724 and XPC-5462 represent a new class of small molecule compounds. These compounds target inhibition of the Na_V_1.6 and Na_V_1.2 channels in excitatory pyramidal neurons and possess a molecular selectivity of >100 fold against Na_V_1.1 channels that are dominant in inhibitory cells. This profile will enable pharmacological dissection of the physiological roles of Na_V_1.2 and Na_V_1.6 and help to define the role of each channel in disease states. These compounds bind to and stabilize the inactivated-state of the channels, demonstrate higher potency with longer residency times, and slower off-rates than carbamazepine and phenytoin. These compounds possess cellular selectivity ex vivo in inhibiting action potential firing in cortical excitatory pyramidal neurons, whilst sparing fast spiking inhibitory interneurons. XPC-5462 also suppresses epileptiform activity in an ex vivo brain slice seizure model. This class of compounds provides a unique approach for treating neuronal excitability disorders by selectively down-regulating excitatory circuits.

**Graphical Abstract:** 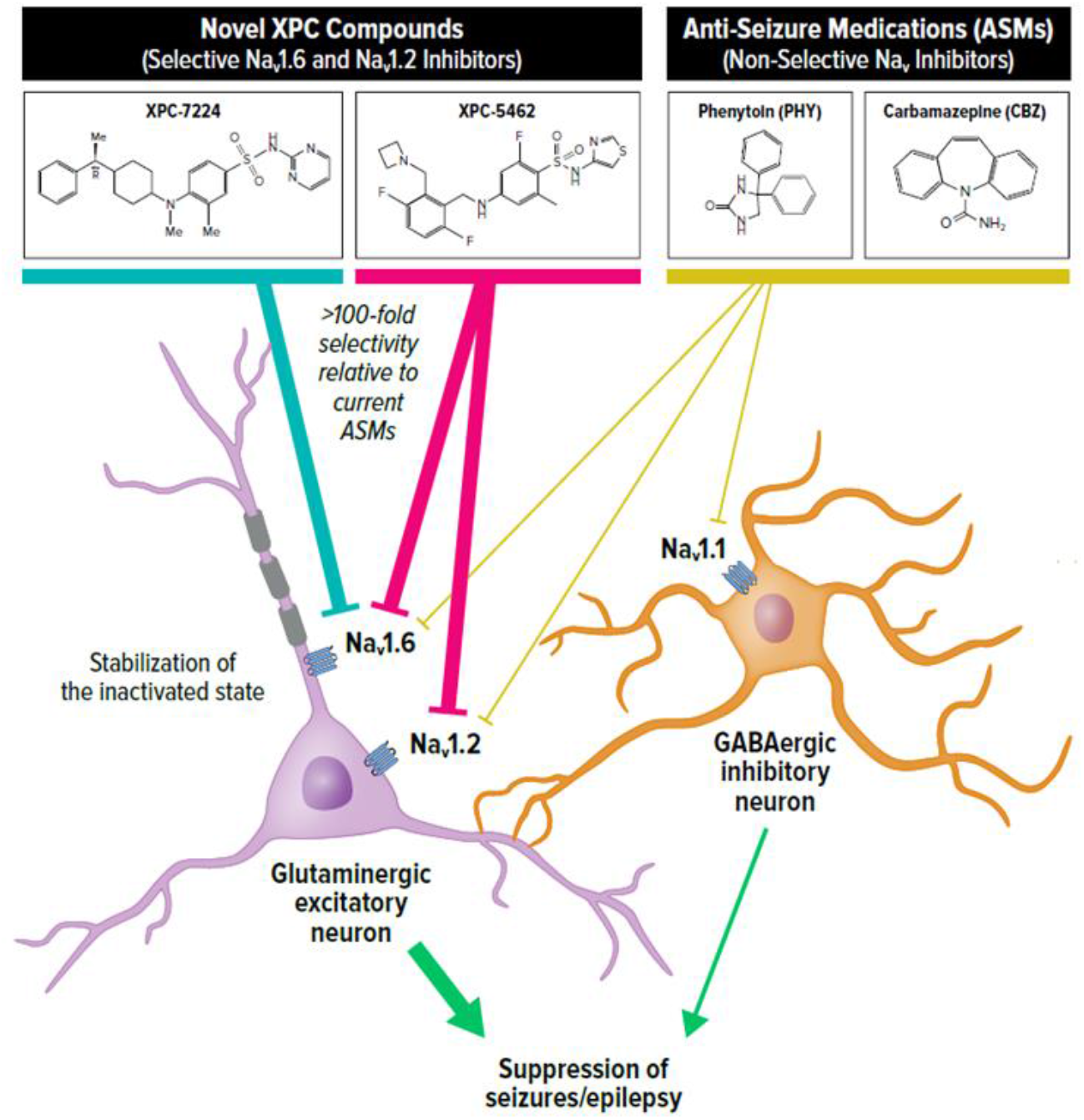

## Introduction

In the central nervous system (CNS), voltage gated sodium channels (Na_V_) control initiation and propagation of action potentials (AP) in excitable tissues.^1-3^ These properties establish Na_V_ channels as attractive pharmacological targets for disorders of hyperexcitability such as epilepsy and pain. Here we introduce new pharmacological tools that enable a dissection of the function of Na_V_1.6 vs Na_V_1.2 and will help to define the role of each channel type in a variety of disease states. We have previously reported the anti-seizure activity of a selective inhibitor of NaV1.6.^4^ Like Na_V_1.6, Na_V_1.2 is also preferentially expressed in excitatory circuits in the CNS. In this report, we introduce 2 pharmacological agents that are very similar in inhibition of Na_V_1.6, but dramatically different as inhibitors of Na_V_1.2. This will enable in vivo studies that evaluate the impact of Na_V_1.2 inhibition.

In the adult CNS, four Na_V_ subtypes are highly expressed: Na_V_1.1, Na_V_1.2, Na_V_1.3 and Na_V_1.6.^5^ Na_V_1.2 and Na_V_1.6 are the major subtypes found in excitatory neurons, whereas Na_V_1.1 is predominantly expressed in inhibitory interneurons.^6^ Several lines of evidence from mouse and human studies now support the parsing of Na_V_ subtypes into excitatory and inhibitory networks, including a many studies indicating that gain-of-function (GOF) in Na_V_1.2 and Na_V_1.6 and loss-of-function (LOF) in Na_V_1.1 are both linked to epilepsy, a disorder of hyperexcitability.^6-15^ The selective expression of Na_V_ subtypes in excitatory and inhibitory circuits rationalizes a selective pharmacological approach in disorders of excitability that targets the excitatory networks whilst sparing the inhibitory networks.

Na_V_ channels are hetero-multimeric proteins composed of large ion conducting α-subunits and smaller auxiliary β-subunits.^16^ The α-subunit is made up of a single transcript that encodes four 6-transmembrane segment domains. Each of these four structural domains can be divided into two functional sub-domains known as the voltage-sensing domain (VSD) and the pore domain (PD). These two functional domains are connected through the intracellular S4-S5 linker to control gating. Na_V_ channels cycle through three basic gating states: rest, open, and inactivated. During depolarizations of sufficient magnitude, outward movement of VSDI-III translate changes in transmembrane potential into channel activation through an electromechanical coupling process to the activation gate at the inner pore.^17-19^ The channels then rapidly enter a fast inactivated state, mediated through the outward movement of VSD-IV. VSD-IV is specialized in its function in controlling inactivation with a hyperpolarized voltage-dependence and a slightly slower transition rate to the UP state than VSDI-III and has been shown to be necessary and sufficient for fast inactivation.^17,20^ The hyperpolarized voltage-dependence of VSD-IV compared to VSDI-III also enables the channels to enter inactivated states without channels opening, known as closed or steady-state inactivation.^21^ Inactivation is one of the mechanisms by which channel availability varies across different resting membrane potentials between neuronal types and sub-compartments of the cell and during neuronal activity.^22^ S4 movement during inactivation triggers an interaction of the hydrophobic motif (IFM) in the DIII-IV linker with another region of the channel.^23^ Structural studies have established the basis of inactivation as an allosteric mechanism where the IFM motif binds into the space between S6 and S4-S5 and translates into a constriction that closes the permeation pathway.^24,25^ Importantly, at typical CNS neuronal resting membrane potentials (RMP) of around -50 mV to -90 mV the channels will be distributed between resting and inactivated states as RMP is often close to the voltage at which channels are equally distributed (inactivation V_0.5_).^26^

Most small molecules that are known to inhibit Na_V_ channels have been non-selective among the isoforms, a function of the pore binding site where key residues are highly conserved across the paralogues.^27-31^ These molecules have found wide ranging clinical efficacy in disorders of excitability including pain, arrhythmia, and epilepsy. Compounds such as Carbamazepine (CBZ) and Phenytoin (PHY), however, have been associated with narrow safety margins. Side effects can appear at doses that overlap with therapeutic doses and may result from over-inhibition of Na_V_ currents and/or off-target interactions.^32-34^ The mechanism by which Na_V_ channels are inhibited by these molecules is multifactorial and includes physical occlusion of the ion conductance,^29^ stabilization of inactivated states,^30^ and immobilization of voltage-sensors.^35^

More recently a new class of selective Na_V_ inhibiting small molecules were identified that interact with extracellular part of VSD-IV and exploit the sequence diversity among the paralogues at this site.^36^ These compounds bind to the activated state of VSD-IV and make an important electrostatic interaction with the fourth arginine residue on VSD-IV-S4 that traps the VSD-IV in the UP state and thus maintains the channel in a non-conducting, inactivated state.^37,38^ Initial compounds of this type potently inhibited Na_V_1.2, Na_V_1.6 and Na_V_1.7 and were highly selective against Na_V_1.1 and Na_V_1.5. Moreover, these compounds had very low CNS penetration. We have now discovered a new CNS penetrant class of VSD-IV targeting compounds with an unprecedented selectivity profile, targeting Na_V_1.6 channels alone,^4^ or Na_V_1.6 and Na_V_1.2 simultaneously for use in epilepsy.^39^ Rodent seizure efficacy data indicates a greater therapeutic index results from Na_V_1.6 selective inhibition compared with the non-selective anti-seizure medications (ASMs), CBZ and PHY, supporting the potential of increased safety margins for selective compounds.^4^

Here, we describe the comparative molecular pharmacology of the Na_V_1.6 selective XPC-7224 and the dual inhibitor of Na_V_1.6 and Na_V_1.2, XPC-5462, to the clinically used non-selective Na_V_ blocking ASMs PHY and CBZ. We characterize several important biophysical and pharmacological differences between these compounds. The XPC compounds bind distinct sites of the channel, have exquisite selectivity for excitatory Na_V_ subtypes, and display radically different kinetic profiles and state dependence of potency when compared with CBZ and PHY. The proposed mechanism for seizure reduction of this pharmacological profile is then demonstrated in patch-clamp recordings from excitatory and inhibitory neurons in adult mice and multi electrode array (MEA) seizure model in acute brain slices. By exploring the differences in selectivity, kinetics, and state dependence, we propose that steady-state inhibition from resting membrane potentials is mechanistically sufficient to reduce excitability in CNS neurons and suppressing seizure-like activity in brain slices.

## Results

### XPC-7224 & XPC-5462 are molecularly selective inhibitors of Na_V_1.6 & Na_V_1.6/1.2, respectively

State-dependence is a common feature of small molecule Na_V_ channel inhibitors, where compounds interact more strongly with the inactivated state of the channel.^30,40^ For instance, traditional local anesthetics target the inner pore cavity to inhibit inactivated or open states of the channel with a greater potency than with closed/rested states. This is also true for ASMs, such as PHY and CBZ (**Figure 1A**), that target the same binding site.^27,41^ The increased potency is proposed to result from a favorable structural configuration in the pore cavity in the inactivated state that increases binding affinity.^42^

**Figure 1:**
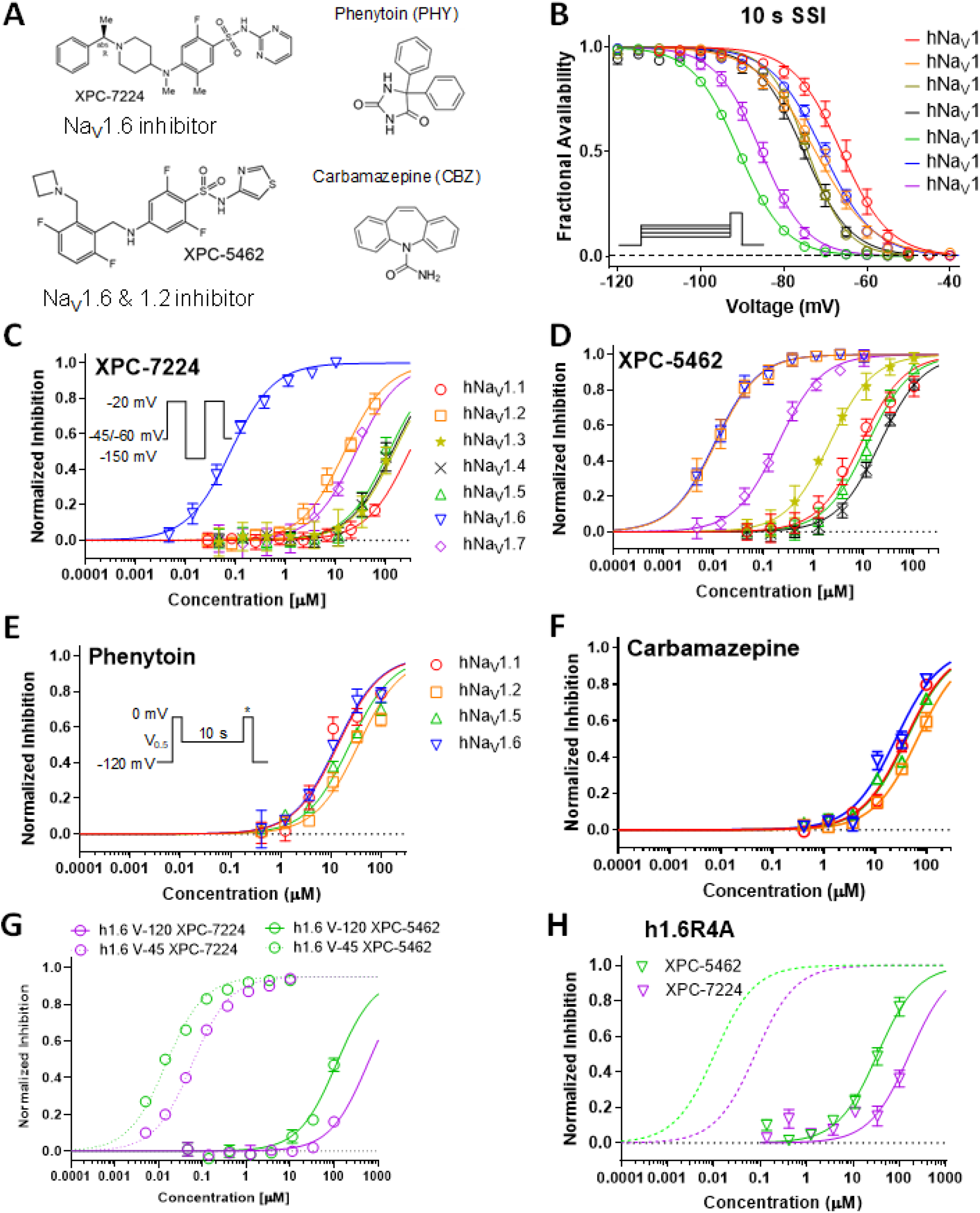
Comparative pharmacology of molecularly selective state dependent inhibitors compared with non-selective ASMs. (A) Structures of XPC compounds and classic pore blocking ASMs PHY and CBZ. (B)Normalized inactivation curves showing different voltage dependence of inactivation across human Na_V_ subtypes. (C)Potency of XPC-7224 plotted as fractional inhibition of different channel subtypes and fitted with a Hill equation. (D)Potency of XPC-5462. (E - F) Potency of the pore blocking ASMs measured using a protocol to capture compounds with fast off rates. (G) Membrane holding voltage dependence of potency. (H) Potencies for h1.6R4A channels.

Selective Na_V_ channel VSD-IV targeting compounds are highly state-dependent, where high affinity binding is dependent upon an interaction with the UP state of VSD-IV that controls the inactivated state.^37^ For accurate measurement of subtype molecular selectivity in voltage-clamp experiments, it is vital to control the channel’s state and apply a transmembrane voltage that ensures equivalent state occupancy for compound binding. To establish this, we assessed the steady-state availability of the subtypes and found V_0.5_ and slope parameters (**Figure 1B** and **Table S1**). Next, concentration responses were measured for the compounds XPC-7224, XPC-5462, PHY, and CBZ at holding-potentials where channels were maintained at fully inactivated potentials based on the fractional availability of the Na_V_ channels (**Figure 1B**). The selectivity profile of the Na_V_1.6 compound XPC-7224 indicates that the IC_50_ for Na_V_1.6 was 0.078 μM (95% CI 0.072 to 0.085 μM), which was >100-fold more potent than all other Na_V_ subtypes tested (**Figure 1C**). The inhibitory selectivity profile for a distinct chemical series, exemplified by XPC-5462, demonstrates equipotent inhibition of Na_V_1.2 and Na_V_1.6 with IC_50_ values in the low nanomolar range (Na_V_1.2 IC_50_ = 0.0109 μM [95% CI 0.00968 to 0.0122 μM] and Na_V_1.6 IC_50_ = 0.0103 μM [95% CI 0.00921 to 0.0115 μM]) (**Figure 1D and Table 1**).

**Table 1:** Potency of Na_V_ channel subtype inhibition of compounds.

|  | XPC-7724 |  |  | XPC-5462 |  |  | Phenytoin |  |  | Carbamazepine |  |  |
| --- | --- | --- | --- | --- | --- | --- | --- | --- | --- | --- | --- | --- |
| | $IC_{50}$ | 95% CI | N (Cells) | $IC_{50}$ | 95% CI | N (Cells) | $IC_{50}$ | 95% CI | N (Cells) | $IC_{50}$ | 95% CI | N (Cells) |
| hNav1.1 | 294 | 255 to 354 | 71 | 8.75 | 7.53 to 10.2 | 62 | 13.8 | 9.59 to 20.1 | 26 | 39.3 | 33.3 to 46.7 | 41 |
| hNav1.2 | 14.9 | 13.0 to 17.1 | 36 | 0.0109 | 0.00968 to 0.0122 | 49 | 35.2 | 2.1 to 44.3 | 26 | 67.4 | 57.3 to 79.6 | 35 |
| hNav1.3 | 147 | 113 to 198 | 47 | 206 | 1.85 to 2.30 | 63 |  |  |  |  |  |  |
| hNav1.4 | 133 | 114 to 158 | 43 | 20.8 | 19.1 to 22.7 | 42 |  |  |  |  |  |  |
| hNav1.5 | 114 | 94.2 to 141 | 23 | 11.9 | 10.5 to 13.5 | 29 | 23.8 | 19.2 to 29.6 | 23 | 41.1 | 35.3 to 47.8 | 39 |
| hNav1.6 | 0.078 | 0.0718 to 0.0848 | 70 | 0.0103 | 0.00921 to 0.0115 | 50 | 13 | 10.1 to 16.9 | 29 | 25.6 | 21.1 to 30.8 | 45 |
| hNav1.7 | 25.3 | 20.0 to 32.8 | 47 | 0.192 | 0.175 to 0.211 | 54 |  |  |  |  |  |  |
| mNav1.6 | 0.13 | 0.103 to 0.164 | 30 | 0.0137 | 0.0112 to 0.0169 | 30 |  |  |  |  |  |  |

The potency of these compounds on mouse Na_V_1.6 channels was also found to be very similar to human channels suggesting no orthologue difference (**Figure S1 and Table 1**). As it was not possible to detect potent inhibition when using the fully inactivated state protocol, pore targeting ASMs were tested by holding cells at the empirically determined inactivation V_0.5_ potential for each cell for 10 s before a test-pulse. Holding the cells at V_0.5_ also ensured that the state occupancy is equivalent in each subtype tested allowing the determination of molecular selectivity. This suggests that the unbinding rates of these compounds at -150 mV are significantly faster than the recovery rate from inactivation. PHY and CBZ were much lower in potency and effectively non-selective with IC_50_ values within 3-fold for CNS channels and Na_V_1.5 (**Figure 1E and F, Table 1**). These results establish the molecular selectivity and increased potency of this new class of XPC compounds relative to traditional Na_V_ targeting ASMs.

### XPC-7224 & XPC-5462 potencies are highly inactivated state-dependent

We next tested the XPC compounds’ potency at a membrane potential of -120 mV, where the channels were fully available at rest. The potency for Na_V_1.6 was reduced by >1000-fold from 0.078 μM to >100 μM for XPC-7224 (n=31 cells) and from 0.010 μM to >100 μM (n=34 cells) for XPC-5462, indicating very strong preference for interacting with the inactivated state of the channel (**Figure 1G**).

### High potency inhibition by XPC-7224 & XPC-5462 depends on the 4^th^ positively charged residue on the DIV-S4 (R1626) in hNa_V_1.6

A previous study using X-ray crystallography in Na_V_1.7 determined that an aryl sulfonamide (GX-674) compound binds the VSD-IV segment.^37^ This interaction between VSD-IV and the compound is mediated via the negatively-charged sulfonamide group of GX-674 and the fourth positively charged arginine side chain (R4) on VSD-IV-S4 (R1608).^37^ Functional studies also demonstrated that neutralizing the R4 charge with alanine mutagenesis (h1.7-R1608A) led to loss of high-affinity binding.^36,37^ These XPC selective inhibitors are also aryl sulfonamides (**Figure 1A**) but differ substantially in chemical scaffold to the previously developed Na_V_1.7 targeting compounds. To test if the corresponding Na_V_1.6 VSD-IV-S4 arginine side chain (R1626) comprises part of the high-affinity binding site for these compounds, we tested Na_V_1.6 (R1626A). The inactivation gating of the mutant was found to be more stable with a ∼20 mV left-shifted V_0.5_ compared with WT (**Figure S2**). The R1626A point mutation resulted in a >1000-fold decrease in potency with an IC_50_ of 166 μM (95% CI 120 to 239 μM, n=29 cells) and 33.6 μM (95% CI 27.1 to 41.8 μM, n=34 cells) for XPC-7224 and XPC-5462, respectively (**Figure 1H**). This positively charged residue on hNa_V_1.6 VSD-IV-S4, therefore, comprises a critical part of the high-affinity binding interaction for these compounds. These findings highlight the similarities between XPC compounds and the GX-674 interaction with Na_V_1.7, establishing the molecular basis for the strong state-dependent binding of XPC-7224 and XPC-5462 to hNa_V_1.6.

### XPC-7224 & XPC-5462 delay recovery from inactivation

Previous studies have established that recovery from inactivation must occur before channels can re-open and the time-dependence of this process reflects the return of VSD-IV to the down state.^43^ We measured the concentration-dependence of the recovery from inactivation of Na_V_1.6 in the presence of the compounds and found that the compounds introduced a slow component for channel recovery (T_slow_) (**Figure 2A-D**). XPC-7224 and XPC-546 had a T_slow_ of ∼20 s, PHY had a T_slow_ of ∼3 s, while CBZ did not exhibit a slower component in the recovery rate. The T_slow_ fraction of recovery increased as the compound concentration increased. **Figure 2E** shows a plot of the concentration-dependence of T_slow_ and indicates that it did not vary significantly with concentration. As ligand unbinding rate (k_off_) is independent of concentration, these data support that T_slow_ reflects the unbinding rate of the compounds at -120 mV, which is rate-limiting the return of VSD-IV back to the resting state. As no change was detected for CBZ, this suggests that the unbinding rate for this compound was much faster than the rate of recovery of the channel from inactivation.

**Figure 2:**
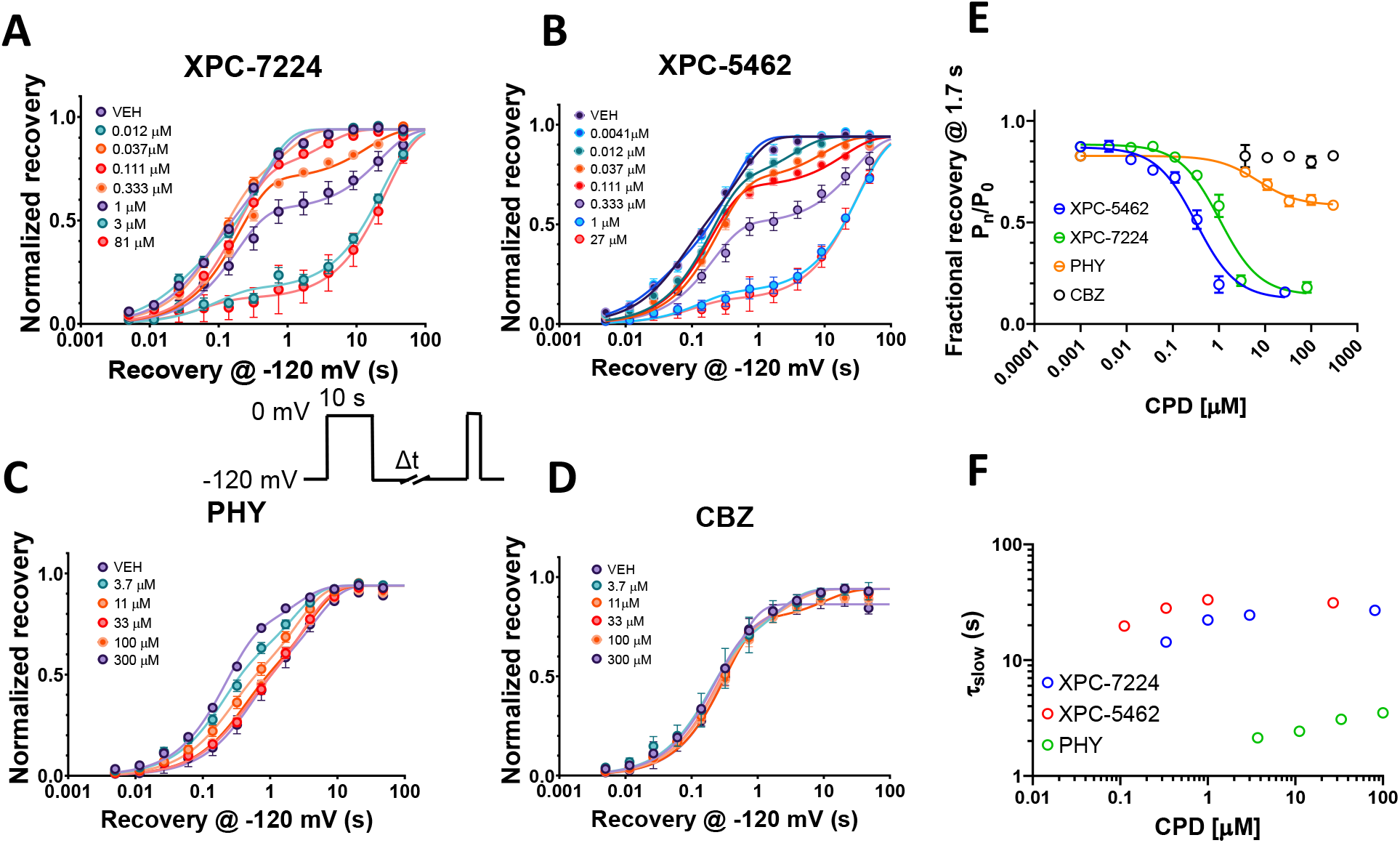
Recovery from inactivation is limited by compound unbinding. Plots of normalized recovery from inactivation induced by a 10 s pre-pulse to 0 mV in the presence of different concentrations of compounds at -120 mV measured using the protocol shown in inset. (A) XPC-7224, n=4-12 cells per concentration (B) XPC5462, n=3-8 cells per concentration (C) PHY, n=6-17 cells per concentration (D) CBZ, n=3-13 cells per concentration. (E) Time constants of the slow component of recovery induced by compound binding against concentration of compound measured at 1.7 seconds. (F) Plot of the fractional recovery after 1.7 seconds against compound concentration. Mean data were fit with a double exponential function.

We quantified the concentration-dependence of the fraction T_slow_ by using the recovery at 1.7 s, a time-point that best separates T_fast_ and T_slow_ where unbound channels have recovered and bound ones have not (**Figure 2F**). This representation of the fraction of T_slow_ varied with concentration and was analyzed with the Hill equation to give IC_50_ values of 1.04 μM (95% CI 0.836 to 1.29 μM, n=78 cells) and 0.322 μM (95% CI 0.252 to 0.413 μM, n=63 cells) for XPC-7224 and XPC-7462, respectively, and 6.94 μM (95% CI 3.10 to 14.2 μM, n=109 cells) for PHY. Interestingly, these IC_50_ values are lower than the potency for the fully inactivated-state shown in Error! Reference source not found. and **Figure 1** for XPC-7224 and XPC-5462, but they are similar to the PHY IC_50_. These data indicate that XPC compounds require more than 10 seconds to fully equilibrate with the inactivated states of the channel, whereas PHY equilibrated within 10 seconds. This suggests that XPC compounds have slower binding kinetics than PHY.

### XPC-7224 and XPC-5462 have slower binding kinetics compared with PHY and CBZ

Equilibrium IC_50_ measurements are commonly used as a proxy for the dissociation constant (K_d_), assuming that inhibition is tightly coupled to binding. Equilibrium measurements, however, yield no information on the kinetics of binding, which may play an important role in the physiological activity of a drug and are determined by intrinsic chemical rates of association and dissociation (k_on_ and k_off_). k_on_ and k_off_ are proportional to the transitional state energy barriers encountered by a ligand when engaging or dissociating from the binding site and, therefore, give information about what is driving potency; the rate of binding or unbinding or a combination of both.^3^ We measured the kinetics of binding to Na_V_1.6 channels using a protocol that tracks the inhibition over 100 s holding at the V_0.5_ (**Figure 3A**). The potencies for the compounds are measured at the end of the 100 s at the V_0.5_ (**Figure 3B**). The mean normalized rate of inhibition was observed at approximately 3X the IC_50_ of each compound to enable comparison of the kinetics at a concentration that is proportional to the equilibrium IC_50_ (**Figure 3C**). The mean data were fit with a single exponential function to give the following T_obs_ values; XPC-5462: 36.8 s (95% CI 34.9 to 38.8 s, n=69 cells), XPC-7224: 84 s (95% CI 62.1 to 125 s, n=65 cells), CBZ: 0.29 s (95% CI 0.182 to 0.419 s, n=52 cells) and PHY: 1.86 s (95% CI 1.67 to 2.06 s, n=56 cells). CBZ and PHY are >100X and >∼20X faster, respectively, to equilibrate than XPC compounds, highlighting a profound kinetic distinction between the mechanisms of action of traditional pore-blockers and our XPC compounds.

**Figure 3:**
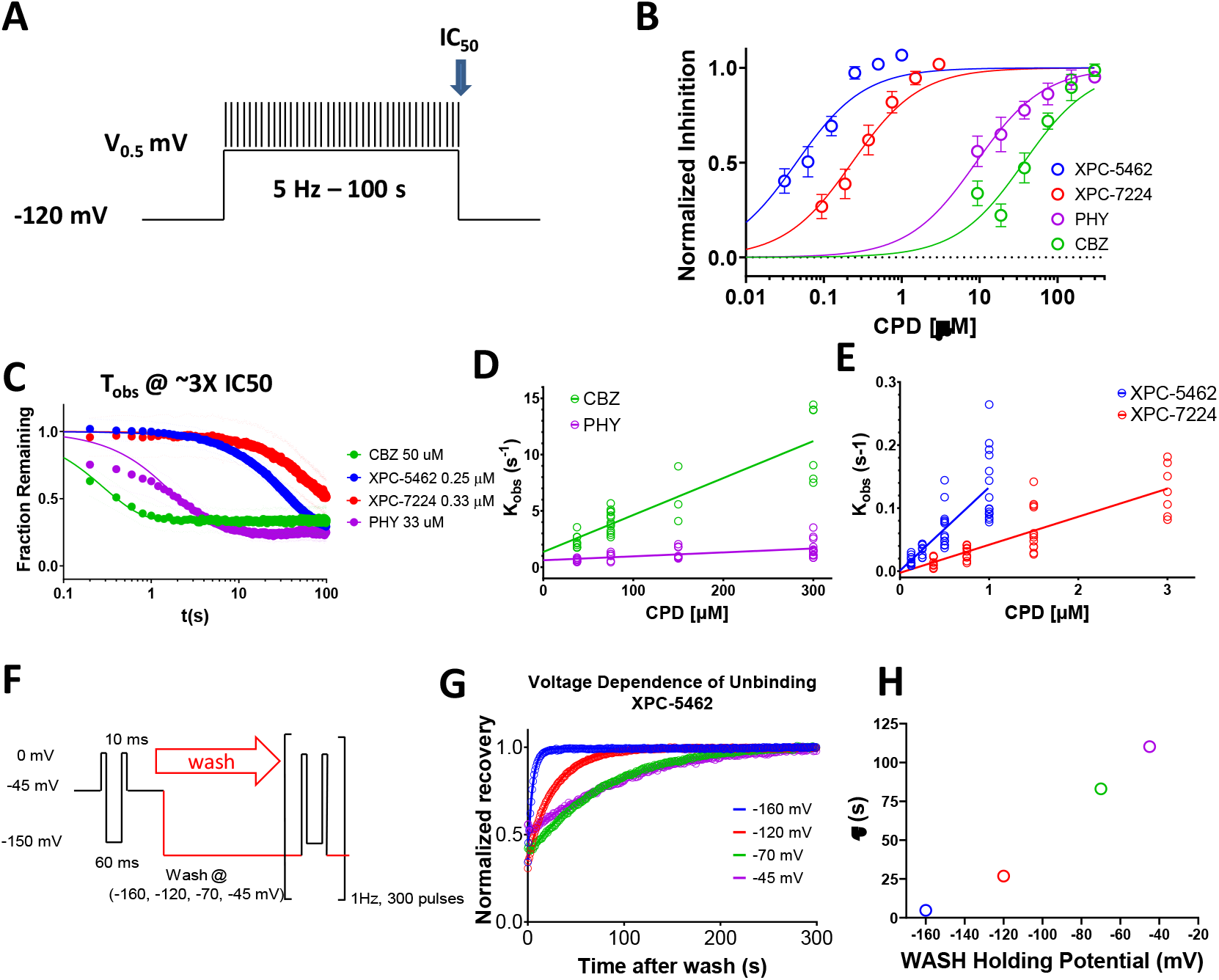
VSD-IV Binding XPC compounds display slower kinetics than typical pore binding ASMs. (A) Protocol used to establish kinetics of compound equilibration at a V_0.5_ holding potential where voltage is stepped to the V_0.5_ for 100 s whilst applying a test pulse at 5 Hz to assess the rate of inhibition. (B) Plot of normalized concentration response fit with the Hill equation to calculate IC_50_ values for Na_V_1.6 inhibition for XPC-5462 (n=7 cells), XPC-7224 (n=3 cells), PHY (n=4 cells) and CBZ (n=3 cells). (C) Normalized equilibration rates of compounds at approximately 3X the IC_50_ concentrations, dots represent 95% CI. (D – E) Plot of K_obs_ against concentration fit with an equation for a straight line to give k_on_. (F) Protocol used to assess the voltage dependence of the unbinding rate in which the compound is equilibrated with the CPD in the inactivated state at -45 mV before simultaneously washing the CPD whilst stepping the voltage. (G) Plot of normalized fractional recovery against time for different holding voltages fit with single exponential functions. (H) Time constants of the recovery rate plotted against membrane voltage during the wash.

By assessing the T_obs_ over a wider range of increasing concentrations, starting at approximately the IC_50_ (**Figure S3**), the k_on_ for each compound can be estimated.^44^ We found that T_obs_ was concentration-dependent (**Figure 3D and E**), as expected from a pseudo 1^st^ order kinetic reaction scheme in which the rate of relaxation following a concentration change is described by K_obs_ = k_on_[CPD] + k_off_ where [CPD] is the concentration of XPC and K_obs_ = 1/T_obs_. By plotting the K_obs_ against concentration, we obtained fitted parameters for k_on_, which for XPC-5462 was 1.32 × 10^5^ M^-1^s^-1^ (95% CI 1.02 to 1.62 × 10^5^ M^-1^s^-1^), XPC-7224 0.445 × 10^5^ M^-1^s^-1^ (95% CI 0.344 to 0.546 × 10^5^ M^-1^s^-1^), CBZ 0.328 × 10^5^ M^-1^s^-1^ (95% CI 0.263 to 0.393 × 10^5^ M^-1^s^-1^) and PHY 0.0349 × 10^5^ M^-1^s^-1^ (95% CI 0.0183 to 0.0515 × 10^5^ M^-1^s^-1^). Interestingly, the k_on_ rates for CBZ and PHY were similar to or slower than the k_on_ for XPC-7224 and XPC-5462 suggesting the greater potency of XPC compounds is driven by more stable binding and longer residence times (**Figure 3C**). Therefore, the kinetic differences at potency matched concentrations are driven by a combination of slower k_off_ and largely the lower concentrations of XPC compounds which reduces the rate of equilibration by mass action (k_on_*[CPD]).

### Off rates are voltage-dependent and reflect compound unbinding

As membrane voltage is not constant in the neuronal physiological environment, we sought to understand the effect of membrane voltage on k_off_. To assess k_off_ directly, we equilibrated the compounds with the channels at a holding-potential of -45 mV with test-pulses every 10 seconds for 5 min (**Figure 3F**). Next, the membrane voltage was simultaneously clamped to a negative holding potential and compound solution exchanged for control solution. The experiment was executed with XPC-5462 at a concentration close to the IC_50_ value (20 nM), to ensure that the wash step fully cleared all residual compound from the microfluidic chamber housing each cell. The recovery of current after washing with XPC-5462 was normalized to the recovery of control cell currents, which were on the same plate but only exposed to vehicle and had accumulated some inactivation over the -45 mV holding period (**Figure 3G**). The normalized recovery traces were fit with single exponential functions to extract T_off_ at different membrane holding-potentials (**Figure 3H**). The more negative membrane potentials had the effect of accelerating the XPC-5462 unbinding (**Figure 3H**). As the downward force on the VSD-IV-S4 is increased with hyperpolarization, the high affinity binding-site becomes deformed, causing the compound to dissociate.

### Membrane potential modulates the apparent potency of XPC-7224 & XPC-5462 through controlling the state of the VSD-IV

To examine the membrane voltage-dependence of inhibition of Na_V_1.6 over a physiologically accessible voltage range, compound’s apparent potency (IC_50, app_) was measured at voltages where channel inactivation rapidly changes.^41^ Potency was assessed by depolarizing the voltage in a step-wise manner from -90 to -70 with 180s intervals at each voltage (**Figure 4A**). The potency had a V_0.5_ of -76.5 mV (95% CI -77.0 to -75.9 mV) and slope of 5.76 (95% CI 5.06 to 6.54, n=46 cells) (**Figure 4B**). To find the membrane potential-dependence of IC_50, app_ the normalized mean fractional inhibition from the last (180^th^) pulse as a function of compound concentration was fit with a Hill equation. The strong membrane voltage-dependence of XPC-7224 and XPC-5462 potency varied over 20 mV from -90 mV to -70 mV by 100-fold, from a low μM IC_50, app_ to the nM range (**Figure 4C and D, Table S2**). For comparison, the curves for the fully inactivated state potency held at -45 mV is shown in dashed lines for XPC-7224 and XPC-5462. The membrane voltage-dependence of IC_50, app_ for PHY and CBZ is lower with only a 10-fold difference between -90 mV and -70 mV (**Figure 4E and F**).

**Figure 4:**
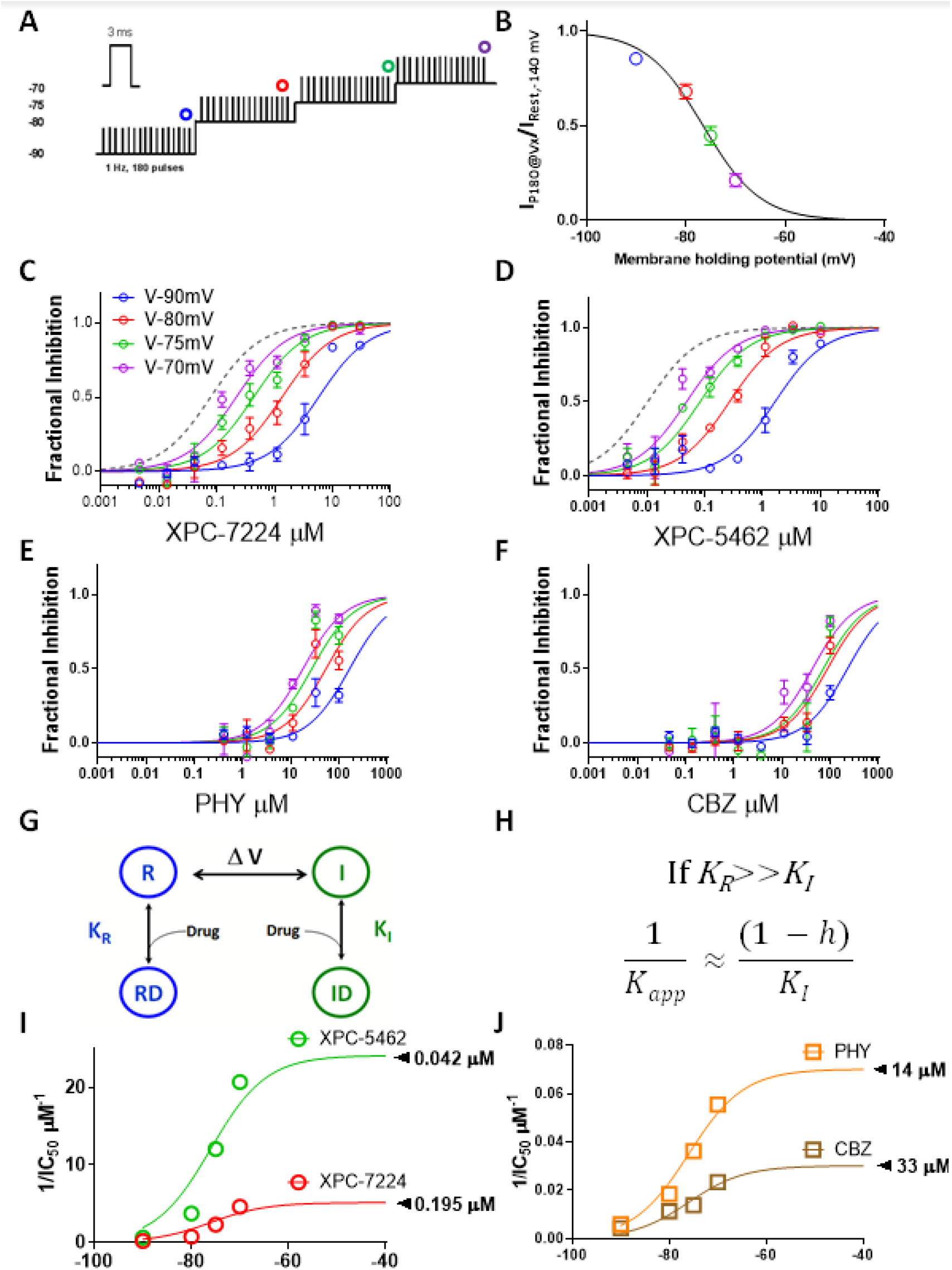
State dependence of hNa_V_1.6 inhibition. (A) Protocol used to assess state dependence of potencies. 180, 3 ms pulses to 0 mV were applied at 1 Hz at each holding potential. (B) Voltage dependence of inactivation in the protocol shown previously. After 3 minutes at each holding potential the availability of channels was measured and normalized to resting state availability and plotted against voltage. (C-F) Normalized fractional inhibition of channels measured at the 180th pulse at different voltages as indicated. In grey dashed line is the curve fit from fully inactivated Na_V_1.6 channels held at -45 mV (Figure 1) (G) 4-state scheme used to model interaction of compounds with inactivated or resting state channels. (H) Model prediction for variation of potency with state occupancy. (I-J) Plots of 1/IC_50_ fitted with the model.

To test if the shift in potency correlated with the fractional availability of the inactivated state, we tested a four-state binding model that assumed binding affinity to the inactivated state was much higher than the resting-state (**Figure 4G-H**).^41^ The model predicts that the steady-state affinity (K_app_) will vary proportionally to the fraction of channels in the inactivated state according to the equation in **Figure 4H** where *h* represents the probability of inactivation curve from **Figure 4B**. The data are well fit by this equation and the projected maximal potencies (**Figure 4I-J**) are close to the inactivated state IC_50_’s previously described (Error! Reference source not found.). These data suggest that the potency of the compounds is directly determined by the proportion of channels in the inactivated-state and indirectly controlled by the membrane voltage.

### XPC-7224 & XPC-5462 inhibit sodium currents independently of high frequency firing

To assess the use-dependence of our VSD-binders, we used a protocol that measured use-dependence from the resting states and from a partially inactivated state (empirically derived V_0.5_), which better represents neuronal RMP (**Figure 5A**). The mean V_0.5_ for all cells was -62.6 ± 0.269 mV, n=177. The normalized current inhibition at 1^st^ (steady resting-state inhibition) and 500^th^ pulse (use-dependent inhibition) in each of the four compounds from the -120 mV holding potential show that there is minimal inhibition at any concentration due to the previously described state-dependence (**Figure 5B**). **Figure 5C** shows the partially inactivated steady-state and use-dependent potency from a V_0.5_ holding-potential. The IC_50_ for steady-state inhibition from a V_0.5_ holding potential (P1) for XPC-7224 was 0.196 μM (95% CI 0.154 to 0.250 μM, n=49 cells); XPC-5462 0.0376 μM (95% CI 0.0305 to 0.0462 μM, n=41 cells), PHY 12.1 μM (95% CI 10.3 to 14.2 μM, n=32 cells) and CBZ 17.0 μM (95% CI 13.4 to 21.7 μM, n=25 cells). For UDB (P500) from V_0.5_ the IC_50_ for XPC-7224 was 0.197 μM (95% CI 0.152 to 0.258 μM); XPC-5462 0.0334 μM (95% CI 0.0261 to 0.0426 μM), PHY 6.80 μM (95% CI 5.67 to 8.14 μM) and CBZ 16.2 μM (95% CI 11.9 to 21.9 μM). These results indicate that neither the pore-blocking PHY nor our VSD-blockers have a large difference in the steady-state versus use-dependent potency from either -120 mV or the V_0.5_. This suggests the VSD-IV targeting XPC compounds might act primarily through interacting with and inhibiting the steady-state rather than increasing inhibition over short high frequency bursts. Given that selective VSD-binders show efficacy in animal models ^4^ these data suggest that use-dependent block is not required for inhibition of hyperexcitability from moderate membrane voltages.

**Figure 5:**
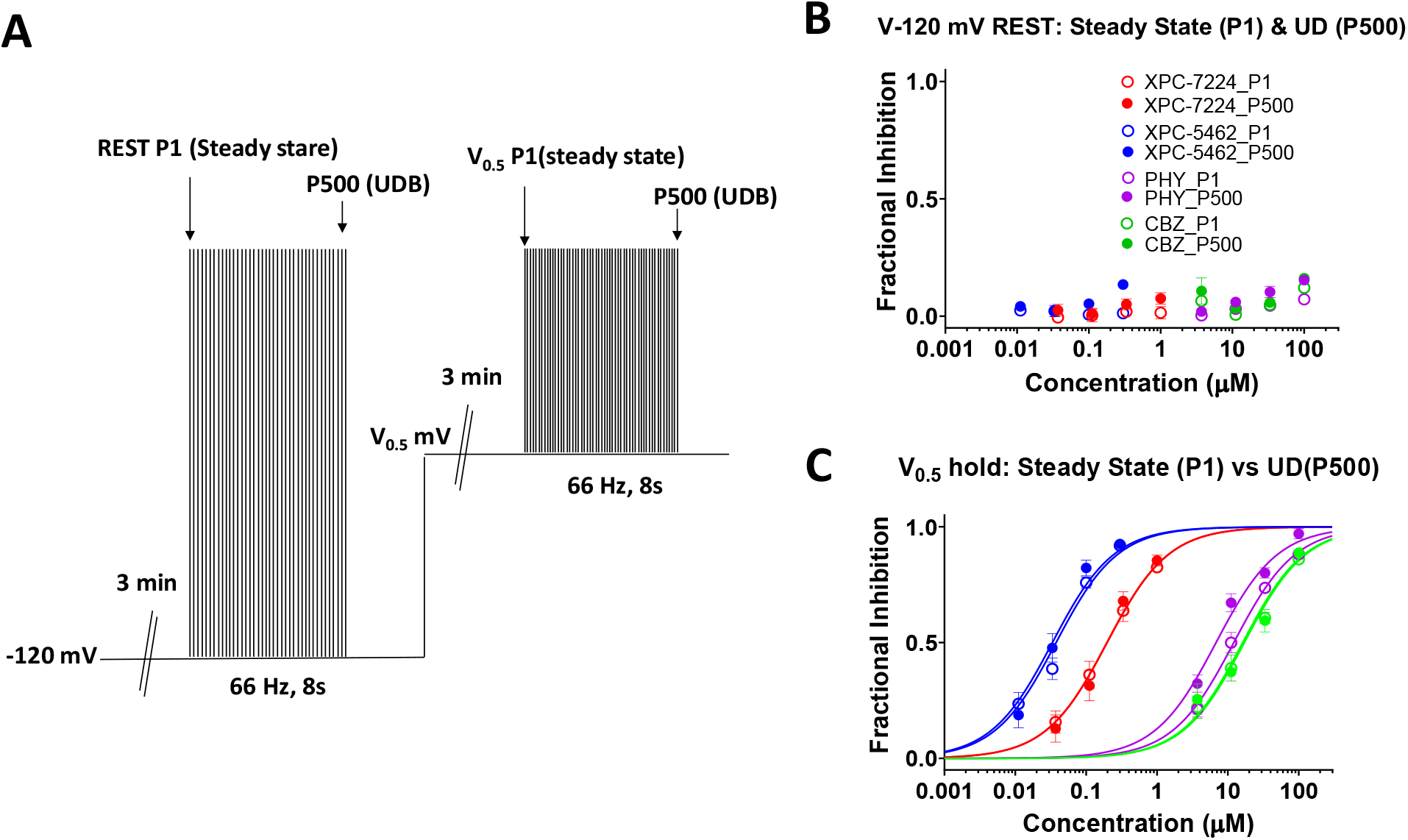
Steady-state inhibition from resting membrane potentials is sufficient for inhibition. (A) Protocol used to assess steady state and use-dependent inhibition from -120 mV or the empirically derived V_0.5_. (B-C) Plot of concentration response of normalized inhibition at either the first pulse (steady state) or the 500th pulse (use-dependent) from a holding-potential of -120 mV (B) or the V_0.5_ (C). n=4-15 cells per concentration data point.

### Inhibiting Na_V_1.6, or Na_V_1.6 and 1.2 selectively suppresses excitatory cortical neurons ex vivo

To test the cellular selectivity of these compounds, we recorded the intrinsic excitability of both excitatory and inhibitory neurons in layer 5 of somatosensory cortex in brain slices. The selective inhibitor of Na_V_1.6, XPC-7224, and the dual inhibitor of Na_V_1.6 and Na_V_1.2, XPC-5462, at 500 nM and 150 nM, respectively, suppressed action potential firing in excitatory neurons, but spared inhibitory interneurons (**Figures 6A-B, D-E**). In contrast, 100 μM of the non-selective pore-targeting inhibitor CBZ inhibited both excitatory and inhibitory cells (**Figures 6C and 6F**). We chose the concentrations of compounds to be approximately 3-fold higher than the potency determined at -70 mV (**Table S2**), which is -77.6 mV with liquid junction potential (LJP) corrected, to approximately match LJP corrected RMP in these neurons (Pyramidal cells RMP was -78.4 ± 2.00 mV, n=13; Fast spiking interneurons RMP was -77.7 ± 1.00 mV, n=10). These data confirm that the Na_V_1.6 and 1.6/1.2 dual targeting molecularly selective inhibitors act as specific excitatory neuron inhibitors.

**Figure 6:**
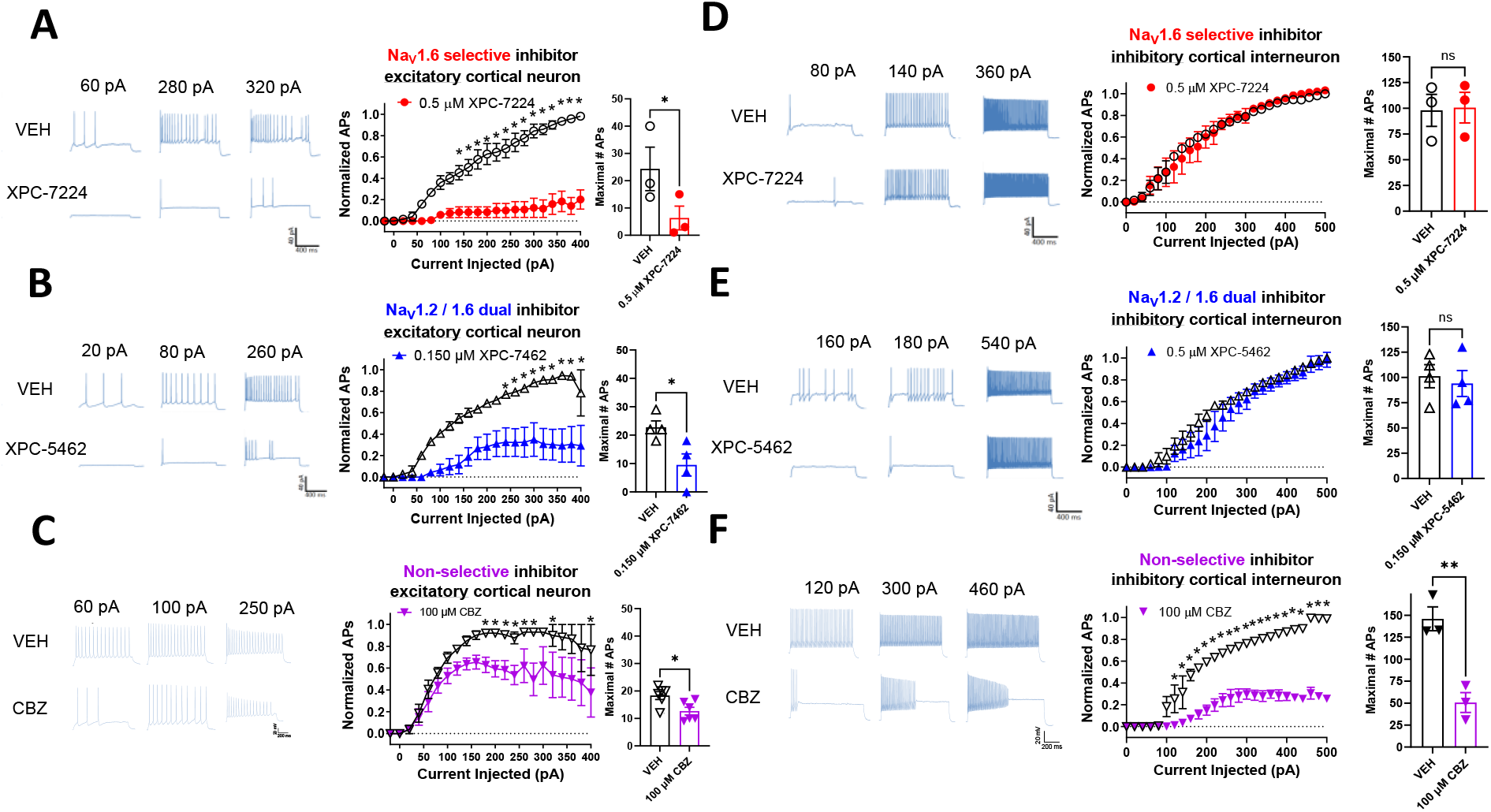
NaV1.6 and NaV1.6 & 1.2 targeting compounds selectively suppress action potentials (APs) in cortical excitatory pyramidal cells. Representative voltage traces from current-clamp recordings of cortical neurons (A-C) and fast spiking interneurons (E-F) from selected depolarizing current injections before and after 10 minutes of incubation with the stated concentration of compounds. Middle panels show input–output plots of the number of APs fired normalized to the maximum number of APs of the vehicle control in the same cell against depolarizing current injection magnitude. Right panels show plots of the absolute maximum number of APs from the range of current injections in vehicle or compound. (A) XPC-7224 at 500 nM significantly inhibited AP firing from pyramidal cells (n=3) but not fast spiking interneurons (n=3) (D). (B) XPC-5462 at 150 nM significantly inhibited AP firing from pyramidal cells (n=4) but not fast spiking interneurons (N=3-4) (E). (C) CBZ at 100 μM significantly inhibited AP firing from pyramidal cells (n=4-6) and fast spiking interneurons (n=3) (B). Statistical significance between vehicle and compound number of APs at each current injection was tested using a 2-way ANOVA followed by Bonferroni multiple comparisons test (*P < 0.05). Students 2-way paired t-tests were used to test the significance of the maximal number of APs fired by each cell in vehicle or compound (\**P*<.05, \*\**P*<.01).

### Suppression of AP firing from CA1 pyramidal cells is modulated by membrane voltage

As XPC-7224 and XPC-5462 are state-dependent inhibitors (**Figure 4**) we hypothesized that they should suppress neuronal excitation more at depolarized membrane potentials. To test this hypothesis ex vivo, we assessed the suppression of AP firing in pyramidal neurons whilst modulating the membrane potential. Both XPC-7224 (500 nM) and XPC-5462 (150 nM) inhibited AP firing elicited by current injection in CA1 pyramidal neurons, as shown in **Figure S4A(i, ii)-B, E(i, ii)-F**, although interestingly with a smaller effect than in neocortical pyramidal neurons (**Figure 6**). The depolarization of membrane voltage decreased AP number, as shown in **Figure S4A(iii), D, E(iii), H**, consistent with our hypothesis that the two compounds stabilized larger fractions of Na_V_ channels in inactivated states upon depolarization.

### Nav1.6 and Nav1.2 suppression suppressed ex vivo seizure-like activity

Finally, we tested the efficacy of these compounds in suppressing neocortical ex vivo seizure-like events using an MEA to assess the effects of selective inhibition of excitatory neurons across an intact network (**Figure S5**). We found that both XPC-5462 and CBZ significantly reduced all epileptiform discharges (**Figure S5E and G**). XPC-7224 had a trending but non-significant effect (P=0.09) on total epileptiform discharges in the 0 Mg^2+^ model (**Figure S5E**), but no clear effect in the 4-AP model. These data demonstrate that isoform specific block of Na_V_1.2 and Na_V_1.6 channels together can significantly limit network seizure-like activity induced by 0 Mg^2+^ or 4-AP.

## Discussion

In this study, we provide a detailed comparative pharmacological and mechanistic characterization of XPC-7224 and XPC-5462, which represent a new class of selective Na_V_ inhibitors. The motivation for developing such a class of compounds was to improve the efficacy and reduce the common adverse event profile of common non-selective Na_V_ inhibiting ASM drugs.^32^ Recently, NBI-921352, a molecule with a similar selectivity profile to XPC-7224 that was created at Xenon, has been shown to be efficacious and have improved tolerability in rodent seizure models when compared with non-selective Na_V_ inhibitors.^4^ Neurocrine Biosciences are currently developing NBI-921352 (Neurocrine, 2019) in phase II clinical trials. These trials will determine the efficacy and safety of selective Na_V_1.6 inhibition in humans as adjunctive therapy to treat SCN8A developmental and epileptic encephalopathy syndrome^45^ and adult focal onset seizures.^46^

Our previous work with NBI-921352 highlighted the selectivity and *in vivo* pharmacological profile of this class of compounds but did not provide a detailed characterization of the biophysics and pharmacology. As XPC-7224 shares the same chemical scaffold as NBI-921352 it is likely that the mechanisms expounded here will be broadly applicable across that chemical scaffold. XPC-5462 has a sufficiently different scaffold to XPC-7224 to impart a different selectivity profile to include Na_V_1.2 activity. As we have shown here, however, the two compounds are almost identical in mechanistic profile aside from their selectivity; they interact with the same binding-site (**Figure 1H**) and are similar in potency (**Figure 1C and D, Table 1**), kinetics (**Figure 3C**), and membrane voltage-dependence (**Figure 4**). These pharmacological similarities suggest that XPC-7224 and XPC-5462 can be considered to have the same molecular mechanism of action.

There are several critical features that differentiate these VSD-IV targeting compounds from the Na_V_ targeting ASMs currently in use. Most strikingly, the molecular selectivity for Na_V_1.6 or dual selectivity for Na_V_1.2 and 1.6 of these compounds position them as a unique class of Na_V_ channel inhibitor that could translate into a differentiated therapy for epilepsy. The >1000-fold selectivity for the cardiac isoform Na_V_1.5 reduces the cardiac risk profile significantly. The sparing of Na_V_1.1 by both compounds will alleviate possible liabilities of suppressing inhibitory interneuron activity. This is highly desirable as loss of Na_V_1.1 conductance is the pathophysiological mechanism driving hyperexcitability in early infantile onset epileptic encephalopathy 6 or Dravet syndrome.^6,47^ In addition to selectivity, the potency range (<200 nM) of these compounds is at least one order of magnitude higher than CBZ, PHY and other putative Na_V_ targeting ASMs such as lamotrigine and lacosamide that are in the 10–100 μM range.^48,49^

The discovery of Na_V_1.6 and Na_V_1.2 targeting compounds was achieved via a progressive medicinal chemistry driven evolution of aryl sulfonamide scaffolds that originally targeted Na_V_1.7 and Na_V_1.3.^39^ The binding mode of early aryl sulfonamides has been structurally elucidated via a co-crystallization of Na_V_ channels with the ligand and found to be a pocket in the extracellular aqueous cleft at the top of the activated-state of VSD-IV, which controls Na_V_ channel inactivation.^37,38^ To assess whether XPC-7224 and XPC-5462 engage the same site as the Na_V_1.7 targeting compounds, we neutralized R1626 in Na_V_1.6, the critical VSD-IV-S4 arginine residue for aryl sulfonamide binding identified from the Nav1.7 crystal structure, to alanine. Na_V_1.6 R1626A displayed a >1000-fold drop in potency for both XPC-7224 and XPC-5462, confirming this key binding interaction was conserved by the new scaffold.

We examined how tightly the XPC compounds were bound to the VSD-IV to stabilize the UP state by measuring the rate of recovery from inactivation at –120 mV in the presence of compound. Recovery from inactivation requires the transitioning of the channel back to the resting-state in which all voltage-sensors are in the down position.^43^ We found that XPC-7224 and XPC-5462 introduced a prominent slower second component of recovery that was not strongly concentration dependent, but the fraction of slowly recovering current was concentration dependent (**Figure 2**). The simplest explanation is that the fraction of channels bound at VSD-IV increased with concentration and that the recovery rate of bound channels reflected the dissociation rate of the compound at -120 mV. Pore-targeting Na_V_ inhibitors also stabilize inactivated states and slow recovery from inactivation.^35,41^ We saw a clearly resolvable slow component of recovery introduced by PHY, but not CBZ, indicating CBZ dissociation rate was faster than recovery from inactivation. The time constants of the T_slow_ and the potency of the fraction of T_slow_ contribution tracked with the potency of the compounds in the order XPC-5462>XPC-7224>PHY. These data demonstrate a striking slowing of dissociation rates and increase in the residency time for the XPC compounds on channels compared with PHY and CBZ.

Equilibration rates are a function of on-rate and concentration as well as off-rates according to K_obs_ = k_on_*[CPD] + k_off_.^3^ A faster k_on_ and/or k_off_, therefore, increases equilibration rates. By comparing the equilibration rates at concentrations 3-fold higher relative to the IC_50_, which are similar to the EC_50_ brain concentrations found to suppress seizures in MES rodent models,^4,50^ we found that these XPC compounds had a much slower equilibration rate than CBZ and PHY (**Figure 3C**). Surprisingly, the XPC compounds had faster k_on_ rates than the pore-binding compounds, despite the slower equilibration rate at 3-fold the IC_50_ (**Figure 3F**). This suggests that a faster k_off_ and mass action are mainly responsible for faster equilibration rates of CBZ and PHY than XPC compounds. We also found the k_off_ was accelerated by hyperpolarization, consistent with the destabilization of compound binding to the VSD-IV UP state (**Figure 3H-J**).

Prior experiments have demonstrated that these compounds have a strong preference for binding the inactivated-state of VSD-IV and that the dissociation rate is dependent on membrane voltage (**Figure 3G**). Consequently, there is a strong dependence of potency on membrane voltage (**Figure 4A-F**). By holding the membrane voltage at increasingly depolarized potentials the potency increased dramatically for the XPC compounds compared with PHY and CBZ. We found that the variation in potency was well fit with a 4-state model where the binding affinity to the inactivated-state was considered much greater than to the resting-state (**Figure 4G-J**), like previous classic studies on Na_V_ inhibitors.^41^ This demonstrates that the relationship between potency and membrane voltage is dependent upon the proportion of channels inactivated rather than a direct voltage-dependence (**Figure 4D**). An important consequence of this dependency is that compound apparent potency will vary dependent upon the distribution of membrane potential in a neuron, channel subtype and subcellular localization of channels.^26,51^

It has been hypothesized that use-dependent block may be a desirable feature of an inhibitor molecule for disorders of hyperexcitability, including epilepsy. The rapid spiking and repeated depolarizations of the membrane that occurs during hyperexcitability favor the inactivated-state of the channel. Interestingly, the XPC compounds did not display a significant increase in potency on Na_V_1.6 after high frequency stimulation when the membrane voltage was held at the V_0.5_ (−70.2 mV with LJP correction) (**Figure 5**). This suggests that, with the XPC compounds, steady-state inhibition was sufficient to allow suppression of hyperexcitability before seizure dependent spiking and depolarization occurs. Indeed, even with the more rapidly equilibrating PHY, inhibition only increases ∼17 % after high frequency firing. This is consistent with the prediction from the 4-state model in **Figure 4** that the steady-state apparent potency can only increase 2-fold when moving from a V_0.5_ potential to a fully inactivated potential, which would give an approximate 20% increase in inhibition at non-saturating concentrations (**Figure 5**). These data suggest use-dependence may not be necessary for efficacy of these Na_V_ targeting ASMs.

We next tested the ability of the compounds to impair intrinsic excitability in the more physiological environment of somatosensory cortex neurons within a brain slice. As previously noted, Na_V_1.6 and Na_V_1.2 are dominantly expressed in excitatory neurons whereas Na_V_1.1 is mainly expressed in inhibitory GABAergic neurons.^6^ Inhibition of either Na_V_1.6 alone with XPC-7224 or Na_V_1.6 and Na_V_1.2 together with XPC-5462 led to reduction of intrinsic excitability of cortical pyramidal cells whilst sparing fast spiking inhibitory neurons (**Figure 6A and B**). This suggests the molecular selectivity of the XPC compounds does confer a cellular selectivity. We propose that this property is highly desirable because it leaves the native inhibitory neuronal system intact to assist with maintaining the excitation-inhibition balance. In contrast, the non-selective Na_V_ ASM CBZ suppressed excitability in both excitatory and inhibitory neurons **(Figure 6C**) which we believe may lead to reduced safety margins.^4^ In addition, we demonstrated increased AP suppression in CA1 pyramidal neurons with depolarization (**Figure S4**) supporting the translatability of our in vitro state-dependence experiments **(Figure 4**). This property would be considered desirable in suppressing hyperexcitability of excitatory neurons receiving excessive depolarizing excitatory input.

To further profile the effects of the compounds we evaluated efficacy in two ex vivo seizure models, where spontaneous seizure like activity is evoked by either removing extracellular Mg^2+^ from the slice or adding 4-AP. In both models, comparable effects were found with XPC-5462 and CBZ providing robust suppression of frequency of seizure like activity and epileptiform discharges **(Figure S5D-G**). Surprisingly, Na_V_1.6 inhibition alone with XPC-7224 did not significantly reduce frequency or discharges, although there was a trend towards reduction of discharges in 4-AP. The differential profile of XPC-7224 and XPC-5462 suggests that in these ex vivo models there may be a requirement for inhibition of Na_V_1.2 channels to suppress seizure like activity. We have previously shown selective Na_V_1.6 inhibition is efficacious at preventing seizures in vivo.^4^ The reason for the lack of effect in ex vivo seizure models is not clear to us and requires further investigation. We speculate that to suppress ex vivo seizure-like activity may require a greater fractional inhibition of the total excitatory cell Na_V_ current, which is made up of both Na_V_1.2 and Na_V_1.6, and Na_V_1.2 has a higher relative abundance.^52^

In conclusion, this study provides a detailed comparative pharmacological characterization of a novel class of selective Na_V_ channel inhibitors. We show that XPC-7224 and XPC-5462 have distinct molecular selectivity profiles targeting either Na_V_1.6 or both Na_V_1.6 and Na_V_1.2. The compounds cause inhibition through the same mechanism of binding to VSD-IV and stabilizing the inactivated-state of the channel, which cannot reset until the compound dissociates. This leads to a strong state, and therefore, membrane potential dependence of potency. The kinetics of these compounds are profoundly different from that of the existing Na_V_ targeting ASMs, CBZ and PHY, with slower equilibration rates driven by slow dissociation and longer residence times on the channels. We found the fractional inhibition of Na_V_ current to be similar at V_0.5_ potentials compared with fully inactivated channels, suggesting that these compounds might exert their activity from steady-state inhibition rather than a use-dependent block. This was confirmed through the inhibition of intrinsic neuronal excitability in brain slices from resting membrane potentials, where only excitatory neurons were inhibited by the XPC compounds. In ex vivo seizure models only XPC-5462 and CBZ demonstrated a strong seizure suppression, indicating the importance of Na_V_1.2 inhibition in those models. It is unclear at this point the translational impact to the clinic of the ex vivo seizure models, but it is important to highlight that selective Na_V_1.6 inhibition has already been demonstrated to be effective in rodent seizure models^4^ and even that Na_V_1.6 inhibition may be the primary driver of in vivo efficacy.^50^ Finally, we propose that this new class of selective Na_V_ targeting compounds will lead to increased clinical efficacy with a higher threshold to adverse side effects because Na_V_1.1 mediated inhibitory neuronal excitability is unperturbed.

## Materials and Methods

### Cell lines

Stably transfected FreeStyle 293 F cells (Thermo Fisher), used for electrophysiology experiments up to passage 30, were maintained in DMEM (Gibco Invitrogen) containing 10% fetal bovine serum, and 0.5 mg/mL Geneticin (G418) at 37°C with 5% CO_2_.

### Animals

Male and female CF-1™ (Charles River) were used in this study (age 3-5 weeks). Mice are housed in individually ventilated cages in 12 h light, 12 h dark lighting regime. Animals received food and water *ad libitum*. All animal handling and experimentation involving animals were conducted following approved protocols according to the guidelines of the Canadian Council on Animal Care (CCAC).

### Electrophysiology

#### Transfecting Cell lines

Stable cell lines were transfected with an expression vector containing the full-length cDNA coding for specific human sodium channel α-subunit using Lipofectamine (Thermo Fisher). The Na_V_1.x stable cell lines and accessory constructs used correspond to the following GenBank accession numbers: Human Na_V_1.1 (NM_006920); human Na_V_1.2 (NM_021007); human Na_V_1.5 (NM_198056); human Na_V_1.6 (NM_014191); mouse Na_V_1.6 (NM_001077499); human Na_V_1.7 (NM_002977); human Na_V_1.3 (NM_0069220). The human Na_V_ β1 subunit (NM_199037) was co-expressed in all cell lines. Human Na_V_1.6 channels were also co-expressed with human FHF2B (NM_033642) to increase functional expression. Human Na_V_1.2 channels were also co-expressed with Contactin 1 (NM_001843) to increase functional expression. Cells were used in automated patch-clamp experiments 24 hours postinduction using Doxycycline (Sigma Aldrich).

### Na_V_ channel automated Qube planar patch-clamp assays

Data was collected using the Qube 384 (Sophion) automated voltage-clamp platform using single hole plates. To measure inactivated state inhibition, the membrane potential was maintained at a voltage where inactivation is complete. For each Na_V_ channel subtype, the V_h_ used to quantify compound inhibition were as follows: Na_V_1.6 (−45 mV), Na_V_1.1 (−45 mV), Na_V_1.2 (−45 mV), Na_V_1.3 (−45 mV), Na_V_1.5 (−60 mV), Na_V_1.7 (−60 mV). The voltage was briefly repolarized to a negative voltage (−150 mV) for 20 milliseconds for Na_V_1.5, Na_V_1.7, Na_V_1.3 or for 60 milliseconds for Na_V_1.1, Na_V_1.2, and Na_V_1.6 to allow recovery from fast inactivation, followed by a test pulse to -20 or 0 mV for 10 milliseconds to quantify the compound inhibition. The repolarization step allows compound-free channels to recover from fast inactivation, but compound-bound channels remain inhibited during the subsequent test step. The Qube is an automated electrophysiology instrument that is blinded to cell selections and experimentation, and selection is performed in a randomized manner. All subsequent data filtering and analysis is performed with automated filters that are applied to the entire dataset from a given Qube run. Appropriate filters for minimum seal resistance were applied (typically >500 MΩ membrane resistance and >3 pF for capacitance), and series resistance was compensated at 100%. The pulse protocols were run at 1 Hz for hNa_V_1.7, hNa_V_1.5, hNa_V_1.3, and hNa_V_1.4 or 0.04 Hz for Na_V_1.6, Na_V_1.1, and Na_V_1.2.

To construct concentration response curves, baseline currents were established after 20 minutes in vehicle (0.5% DMSO). Full inhibition response amplitudes were determined by adding tetrodotoxin (TTX, 300 nM) or tetracaine for Na_V_1.5 (10 μM) to each well at the end of the experiment. Compounds were then exposed at a single concentration for 20 minutes. One-sixth of every experimental plate was dedicated to vehicle-only wells that enabled correction for nonspecific drift (i.e., rundown) of the signal in each experiment. For all channel subtypes, inhibition by the compound reached steady state within 20 minutes of incubation.

The current inhibition values (I_(CPD)_) were normalized to both the vehicle (I_control_) and the full response defined by supramaximal TTX (I_TTX_) or tetracaine (for Na_V_1.5) addition responses according to Equation 1:

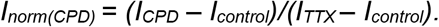

This normalized inhibition was then further normalized to the span of the assay to account for the run-down seen in cells exposed to vehicle alone for 20 minutes as follows:

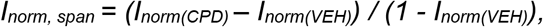

where:

I_*norm, span*_ = the current response normalized to within the span of the assay.

I_*norm*(*CPD*)_ = the normalized response in the presence of compound.

I_*norm*(*VEH*)_ = the normalized response in the absence of compound.

This normalization ensures that the data ranges were between 0 and 1, and there is no rundown in the plots. The normalized data from all cell recordings at a concentration were grouped together and plotted with GraphPad Prism 8, and IC_50_ values were calculated for grouped data using the following version of the Hill equation:

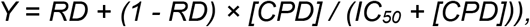

where:

*Y* = the fraction of sodium current blocked in the presence of the compound.

*[CPD]* = the concentration of compound.

*IC*_*50*_ = the IC_50_ concentration.

*RD* = the “rundown” of sodium current in vehicle alone, which is equal to 0 in this case, as the inhibition has already been normalized to the span. The Hill slope was fixed to 1. The normalization of other Qube protocols was performed in the same way. The 95% CI for the IC_50_ from the fitted curve to the mean data were reported and N = the total number of cells that were used in the fit unless otherwise noted.

Recovery from inactivation was measured with a test-pulse to 0 mV following a depolarizing pre-pulse to 0 mV for 10 seconds from a holding potential of -120 mV. The mean normalized recovery of currents from a holding-potential of -120 mV in compound was plotted and fit with a bi-exponential function.

Qube experiments were all performed at 27°C ± 2°C.

### Automated patch-clamp recording solutions

The recording solutions for Na_V_1.1, Na_V_1.2, Na_V_1.3, and Na_V_1.6 cell line studies contained: Intracellular solution (ICS): 5 mM NaCl, 10 mM CsCl, 120 mM CsF, 0.1 mM CaCl_2_, 2 mM MgCl_2_, 10 mM HEPES (4-(2-hydroxyethyl)-1-piperazineethanesulfonic acid buffer), 10 mM EGTA (ethylene glycol tetraacetic acid); adjusted to pH 7.2 with CsOH. Extracellular solution (ECS): 140 mM NaCl, 5 mM KCl, 2 mM CaCl_2_, 1 mM MgCl_2_, 10 mM HEPES; adjusted to pH 7.4 with NaOH. Liquid junction potential (LJP) calculated for these solutions was 7.6 mV and voltages are not corrected unless stated in text. Solutions with a reversed Na^+^ gradient were used for Na_V_1.5 and Na_V_1.7 studies since they improved technical success. ICS: 120 mM NaF, 10 mM CsCl, 0.1 mM CaCl_2_, 2 mM MgCl_2_, 10 mM HEPES, 10 mM EGTA; adjusted to pH 7.2 with CsOH. ECS: 1 mM NaCl, 139 mM CholineCl, 5 mM KCl, 2 mM CaCl_2_, 1 mM MgCl_2_, 10 mM HEPES; adjusted to pH 7.4 with NaOH. Osmolarity in all ICS and ECS solutions was adjusted with glucose to 300 mOsm/kg and 310 mOsm/kg, respectively.

### *Ex v*ivo *electrophysiology*

#### Brain slice patch clamp electrophysiology – cortical recordings

Parasagittal cortical brain slices were prepared from 3-5 week old CF-1 mice from standard procedures adapted from previously published methods.^53^ Briefly, the mouse was deeply anaesthetized with isoflurane and decapitated. The brain was removed and placed into chilled artificial cerebrospinal fluid (aCSF) solution containing (in mM): 125 NaCl, 25 NaHCO_3_, 2.5 KCl, 1.25 NaH_2_PO_4_, 2 CaCl_2_, 2 MgCl_2_, 10 d-glucose, pH 7.3, osmolarity adjusted to ∼306 mOsm using sucrose. All solutions were saturated with 95% O_2_ and 5% CO_2_ constantly perfused with 95% O_2_/5% CO_2_. Parasagittal slices with a thickness of 400 μm were prepared using a vibratome (Ted Pella, Inc.). Following sectioning, the slices were placed in a holding chamber and incubated in a water bath at 34°C for 15 minutes.

Following a 60-minute incubation at room temperature, a brain slice was selected and placed on the stage of an upright microscope (SliceScope Pro 2000, Scientifica). The slice was constantly perfused with room temperature aCSF, containing 0.1% DMSO as a vehicle control, and oxygenated with 95% O_2_/5% CO_2_. The slice was visualized using brightfield microscopy, and a healthy neuron was selected from neocortical layer 5. Whole-cell configuration was achieved with a pipette (bath resistance 4 – 6 MΩ) containing internal solution. Stimulation was applied in current-clamp mode, and consisted of a series of 1000 ms square pulses, beginning at -20 pA and increasing by +20 pA increments (3000 ms between pulses). The internal solution contained (in mM): 120 KMeSO_4_, 20 KCl, 10 HEPES, 4 Na_2_ATP, 2 MgCl_2_, 0.3 Tris-GTP, 0.2 EGTA, pH 7.2-7.3 using KOH or HCl and osmolarity of 285-295 mOsm. LJP calculated for these solutions was +12.7 mV. Experiments were performed at room temp 20-22°C.

### Brain slice patch clamp electrophysiology – CA1 recordings

Brain slices of hippocampal regions were obtained from 3∼5-week-old C57BL/6 male mice, and were prepared with a Vibratome (Leica, VT 1200) at 0 °C. Slicing solution contained (mM): 216 sucrose, 2.5 KCl, 1.25 NaH_2_PO_4_, 10 MgSO_4_, 0.5 CaCl_2_, 11 glucose, 26 NaHCO_3_ (pH 7.2∼7.4; 305∼315 mOsm), constantly bubbled with carbogen. Coronal slices including hippocampal regions were 300 μm thick, prepared with a advancing speed of 0.14 mm/s and a vibration range of 1.2 mm. Slices were then transferred to bath solution (artificial cerebral spinal fluid, aCSF) containing (in mM): 129 NaCl, 2.5 KCl, 1.25 NaH_2_PO_4_, 1 MgSO_4_, 2 CaCl_2_, 10 glucose, 26 NaHCO_3_; 300∼310 mOsm, pH: 7.2-7.4, constantly bubbled with carbogen, and recovered for 30 minutes at 32∼34 °C.

For electrophysiology, glass pipettes were pulled with a P-2000 puller (Sutter Instrument) to generate tips with resistance of 4-6 MΩ. Pipette solution contained (in mM): 128 K-Gluconate, 9 <u>HEPES</u>, 0.5 MgCl_2_, 8 NaCl, 0.1 EGTA, 14 Tris_2_-phosphocreatine, 4 Na_2_-ATP, 0.3 tris-GTP; ∼290 mOsm, pH: 7.2-7.25. Bath solution (aCSF) contained (in mM): 129 NaCl, 2.5 KCl, 1.25 NaH_2_PO4, 1 MgSO_4_, 2 CaCl_2_, 10 glucose, 26 NaHCO_3_; 300∼310 mOsm, pH: 7.2-7.4, constantly bubbled with carbogen. Electrophysiological data were acquired using Multiclamp 700B amplifiers (Molecular Devices) at 32∼34 °C. Data were acquired at 100 kHz and filtered at 50 kHz. Pipette capacitance was compensated by 90% of the fast capacitance measured under Gigahm seal conditions in voltage-clamp prior to establishing a whole-cell configuration. During current clamp, series resistance was < 25 MΩ in all recordings and the bridge was balanced by >70%. Liquid junction potential was corrected after data were obtained. Action potential was counted with a threshold of -20 mV.

Once the recordings in vehicle were completed, and while still holding the patch on the same neuron, the bath solution was changed from 0.1% DMSO in aCSF to compound in aCSF. The slice was incubated in circulating compound for 10 minutes before repeating the series of square pulse stimulations. Working stock solutions were prepared in DMSO at a concentration of 20 mM.

All data analysis was done offline using ClampFit 10.7 (Molecular Devices). Data are presented as a mean ± SEM. For each sweep, the number of evoked APs was counted, and plotted as a function of current injection (beginning with -20 pA) and then normalized to the maximum vehicle response. These generated “input/output” (or “I-O”) curves demonstrating the relationship between stimulus and AP frequency.

### Multi-electrode array (MEA) recordings

Male and female CF-1 mice between 4 to 8 weeks were used in the study. Mice were anesthetized with isoflurane before being euthanatized by cervical dislocation. Brains were then removed and stored in cold cutting solution (in mM): 3 MgCl_2_; 126 NaCl; 26 NaHCO_3_; 3.5 KCl; 1.26 NaH_2_PO_4_; 10 glucose. 350 μm horizontal sections containing somatosensory cortex were made, using a Leica VT1200 vibratome (Nussloch, Germany). Slices were then transferred to a holding chamber and incubated for 1–2 h at room temperature in artificial CSF (ACSF) containing (in mM): 2 CaCl_2_; 1 MgCl_2_; 126 NaCl; 26 NaHCO_3_; 3.5 KCl; 1.26 NaH_2_PO_4_; 10 glucose. All the solutions were bubbled continuously to saturate with carboxygen (95% O_2_ and 5% CO_2_).

MEA recordings were performed on the 3Brain BioCAM DupleX system (Switzerland), using the 3Brain Accura HD-MEA chips with 4,096 electrodes at a pitch of 60μm. Slices were placed onto the electrodes with a harp placed on top to keep the slice pressed down gently to the recording electrodes. Slices were perfused continuously with artificial cerebrospinal fluid (ACSF) and a zero Mg^2+^ ACSF or 100uM 4-Aminopyridine was used to induce epileptiform activity over a 50-minute period. Before removal of Mg^2+^ or addition of 4-Aminopyridine, slices were perfused for 10 minutes in either ACSF with 0.1% DMSO or compound, respectively. Recordings were obtained from the entire slice. Experiments were performed at 33–36°C. The solutions were perfused at the rate of 5.0 mL/min. Signals were sampled at 10kHz with a high-pass filter at 2Hz. Analysis of seizure-like events and epileptiform activity was done using the Xenon LFP Analysis Platform.^54^ Electrophysiological recordings were analysis from the entire neocortical region of the brain slice, sampling every 120 μm. Epileptiform activity was considered seizure-like events when it had a high-frequency rhythmic bursts associated with high-frequency signals lasting at least 10 seconds. All pathological discharges observed from the recordings were considered epileptiform discharges for analysis purposes.

### Quantification and statistical analysis

Quantification of data was done on GraphPad Prism 8 was or ClampFit 10.7 (Molecular Devices) as indicated. Quantitative data are presented as mean ± SEM. The number of cells or brain slices tested are reported as the N in the figure legends where appropriate. Significance was tested using 2-way ANOVA followed by Bonferroni multiple comparisons tests or using students 2-way paired t-tests as indicated in the figure legends.

## Acknowledgments

M-RG was supported by a Mitacs Accelerate fellowship and a Natural Sciences and Engineering Research Council of Canada (NSERC) scholarship. PCR received funding from NSERC. Xenon Pharmaceuticals, Inc employees may hold equity in Xenon Pharmaceuticals, Inc. Neurocrine Biosciences employees may hold equity in Neurocrine Biosciences.

## Author contribution

The experiments in this report were conceptualized by SG, JP, RP, WY, CC. Experiments were conducted and analyzed by SG, AW, NS, MS, RP, ST, WY, and MG. Experiments were also conducted by MW. Cell lines and mutants were created by JM and RD. Medicinal Chemistry expertise was provided by TF. SG, WY, MG, and JP contributed to the writing and editing of the manuscript. All work conducted for this manuscript was under the supervision of PR and JE. All authors approved the final manuscript draft for submission.

