## Supplemental Materials for "Molecular pharmacology of selective Na_v_1.6 and dual Na_v_1.6 and Na_v_1.2 channel inhibitors that suppress excitatory neuronal activity ex vivo"

### Supplementary Information

#### Supplementary Results

##### Suppression of AP firing from CA1 pyramidal cells is modulated by membrane voltage

Depolarization from -80 mV to -60 mV dramatically increases the fraction of $Na_V1.6$ channels (and $Na_V1.2$) in the inactivated conformation (**Figure 1B**), increasing the potency of XPC-7224 and XPC-5462 (**Figure 4C, D**) and reducing the availability of $Na_V1.6$ (and $Na_V1.2$ in the case of XPC-5462) for hyperexcitability. In the presence of the compounds, we injected 60-pA current for 3 minutes, which depolarized cell membrane by ~12 mV from -85.1 ± 1.22 mV to -73.3 ± 1.55 (N=9) (**Figure S4C**) and by ~15 mV from -83.2 ± 1.84 mV to -67.9 ± 2.59 (N=10) (**Figure S4G)**. The depolarization of membrane voltage resulted in a more significant decrease in AP number, as shown in **Figure S4A(iii), D, E(iii), H,** consistent with our hypothesis that the two compounds stabilized larger fractions of $Na_V$ channels in inactivated states upon depolarization. Injection of -150 pA current to hyperpolarize membrane reversed the additional inhibitory effect, as shown in **Figure S4A(iv), D, E(iv), H**, presumably by promoting dissociation of the compounds (**Figure 3I**). These data support the coupling of apparent potency to the state occupancy of the channels in neurons.

##### Inhibiting $Na_V1.6$ or $Na_V1.6$ and 1.2 suppresses seizure-like events in ex vivo model of ictogenesis

The removal of extracellular $Mg^{2+}$ typically results in seizure-like events occurring after approximately 10-20 minutes due to the removal of NMDA receptor block and consequent unrestrained glutamatergic activity.[1] The addition of 4-Aminopyridine is thought to produce seizure like event due to the alteration of interneuron activity by blocking interneuron potassium channels.[1] We used MEA analysis of the electrographic signal over the entire neocortical slice, sampling every 120 µm We used this high-resolution spatial sampling to create raster plots of the local field potential activity over the entire slice to assay epileptiform activity over space and time (**Figure S5Ai-Aiv**). Individual traces are also shown to further demonstrate the type of epileptiform events occurring within the slices (**Figure S5B, C**). Slices bathed in the convulsant media displayed large seizure-like events (**Figure S5Bi, Ci**), however, when slices were bathed in either XPC-5462 or CBZ whilst in the convulsant media, significantly less seizure-like events occurred (**Figure S5B, C, D, F**), indicating that $Na_V1.6$ and $Na_V1.2$ inhibition is sufficient to suppress induced seizure like events. However, short duration epileptiform discharges lasting 1-3 seconds were still present in some channels (**Figure S5Aii, Aiv, Bii, Biv, Cii, Civ**), which are too brief to be classified as seizure-like events (<20 seconds) but are still epileptiform discharges. We found that slices bathed in XPC-7224 whilst also in the convulsant media still presented seizure-like events (**Figure S5Aiii, Biii, Ciii, D, F**) suggesting that in these models, inhibition of $Na_V1.6$ alone is not sufficient to stop seizure like events. We further assayed the total epileptiform discharges from these recordings, to determine if the compounds were also suppressing other pathological activity in addition to the seizure-like events.

#### Supplementary Tables

**Table S1: Steady-state inactivation properties of $Na_V$ subtypes after 10s equilibration at each voltage.**

| | $V_{0.5}$ (mV) | 95% CI | Slope | 95% CI | N (Cells) |
|---|---|---|---|---|---|
| **hNav1.1** | -66.2 | -67.1 to -65.3 | 5.83 | 5.07 to 6.64 | 11 |
| **hNav1.2** | -72.4 | -73.2 to -71.6 | 6.99 | 6.32 to 7.70 | 12 |
| **hNav1.5** | -90.8 | -91.0 to -90.6 | 5.58 | 5.38 to 5.78 | 63 |
| **hNav1.6** | -70.6 | -71.0 to -70.2 | 6.15 | 5.79 to 6.53 | 13 |
| **hNav1.7** | -85.5 | -86.1 to -84.9 | 5.47 | 4.96 to 6.01 | 8 |

**Table S2: Membrane voltage dependence of potencies.**

| | XPC-7224 | | | XPC-5462 | | | Phenytoin | | | Carbamazepine | | |
|---|---|---|---|---|---|---|---|---|---|---|---|---|
| HP | $IC_{50}$ (μM) | 95% CI | N (cells) | $IC_{50}$ (μM) | 95% CI | N (cells) | $IC_{50}$ (μM) | 95% CI | N (cells) | $IC_{50}$ (μM) | 95% CI | N (cells) |
| **-90 mV** | 5.78 | 4.39 to 7.68 | 42 | 1.64 | 1.27 to 2.14 | 38 | 168 | 116 to 256 | 32 | 229 | 169 to 328 | 43 |
| **-80 mV** | 1.34 | 0.979 to 1.85 | 42 | 0.267 | 0.220 to 0.3526 | 38 | 54.2 | 35.1 to 85.1 | 32 | 88.1 | 62.7 to 127 | 43 |
| **-75 mV** | 0.434 | 0.314 to 0.598 | 42 | 0.0829 | 0.0603 to 0.113 | 38 | 27.6 | 18.4 to 42.1 | 32 | 72.1 | 42.7 to 126 | 51 |
| **-70 mV** | 0.217 | 0.154 to 0.309 | 42 | 0.0483 | 0.0345 to 0.0668 | 38 | 18 | 11.1 to 30.4 | 32 | 42.7 | 24.4 to 79.3 | 51 |

#### Supplementary Figures

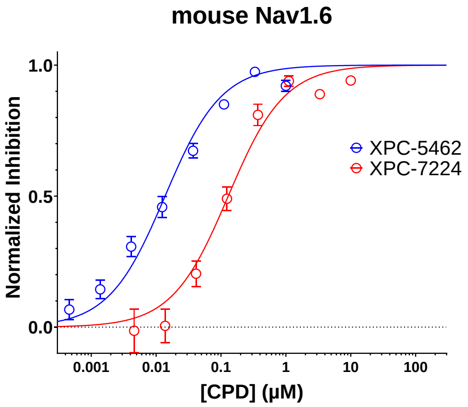


**Figure S1: Potency on mouse Na$_V$1.6.** Plots of mean normalized inhibition at increasing concentrations fit with Hill-Langmuir equation.

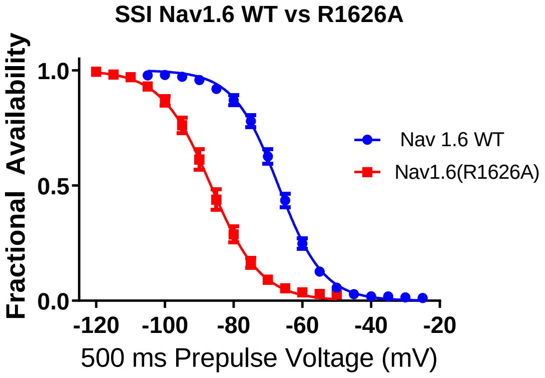


| | Nav 1.6 WT | Nav1.6(R1626A) |
|---|---|---|
| V50 | -66.99 | -86.63 |
| Slope | 6.549 | 7.252 |

**Figure S2: Inactivation gating of Na$_V$1.6(R1626A) is stabilized relative to WT Na$_V$1.6.** Normalized Steady State Inactivation after a 500 ms pre-pulse.

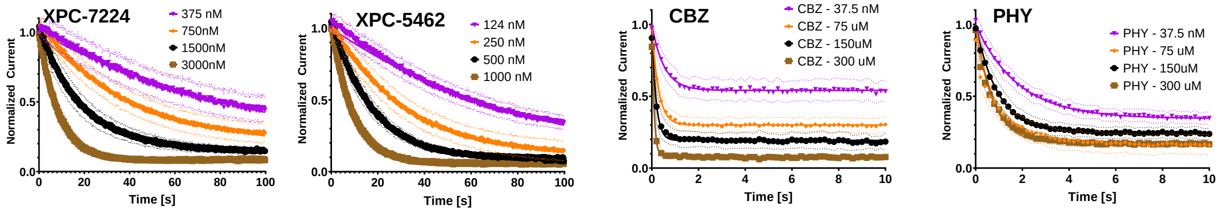


**Figure S3: Concentration dependence of $T_{obs}$.** Plots of mean normalized inhibition at increasing concentrations starting at approximately the $IC_{50}$ fit with single exponential function to give $T_{obs}$.

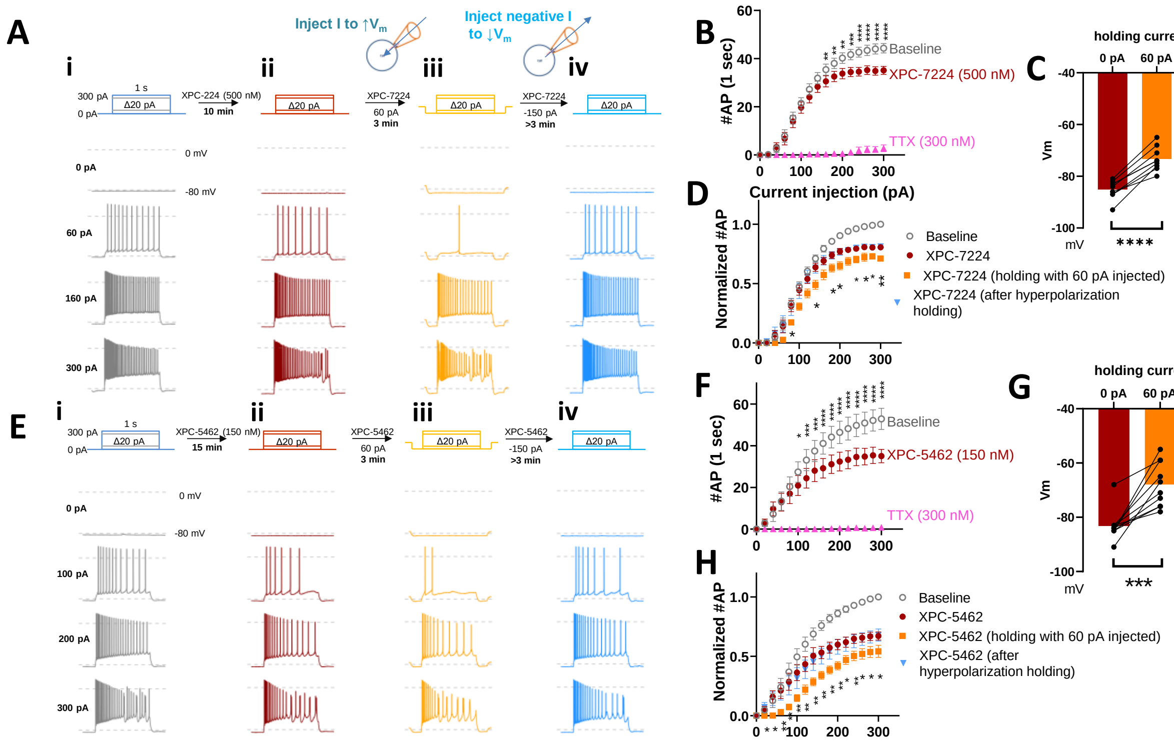

**Figure S4: Suppression of AP firing from CA1 pyramidal cells is modulated by membrane voltage.** (A) Representative traces from current clamp recordings of hippocampal CA1 neurons before and after XPC-7224 (500 nM) application (i and ii), followed by recordings with a protocol spanning the same current-injection range (0~300 pA) with 20-pA increments, but 60-pA current is injected for 3 minutes and at pulse intervals (iii), and at the end recordings after prolonged -150-pA injection for 3~5 minutes (iv). **(B)** Input-output plots showing the number of APs within 1-sec injection against injected depolarizing current magnitude for XPC-7224. Significance was determined with corrected Sidak's multiple comparisons following 2-way ANOVA. TTX, (300 nM) was applied to some cells at the end to confirm perfusion functioning. **(C)** Change of membrane potential in response to injection of 60 pA current. Significance was determined with paired t-test (***P < 0.001, ****P < 0.0001). **(D)** Input-output plots showing the number of APs fired normalized to the maximal number of APs before compound application in the same cell, from recordings as in A (i ~ iv). * show the significance between normalized action potential counts between 0 pA injection and 60 pA injection, determined by Dunnett multiple comparisons following 2-way ANOVA among conditions for XPC-7224. (*P < 0.05, **P < 0.01, ***P < 0.001, ****P < 0.0001). **(E)** Representative traces from current clamp recordings as in (A) for XPC-5462 (150 nM). **(F)** Input-output plots for XPC-5462. Significance was determined as in B. **(G)** Change of membrane potential in response to injection of 60 pA current. Significance was determined as in C. **(H)** Input-output plots showing the number of APs normalized to the maximal number of APs for 150 nM XPC-5462 as in E (i ~ iv). Significance was determined as in D.

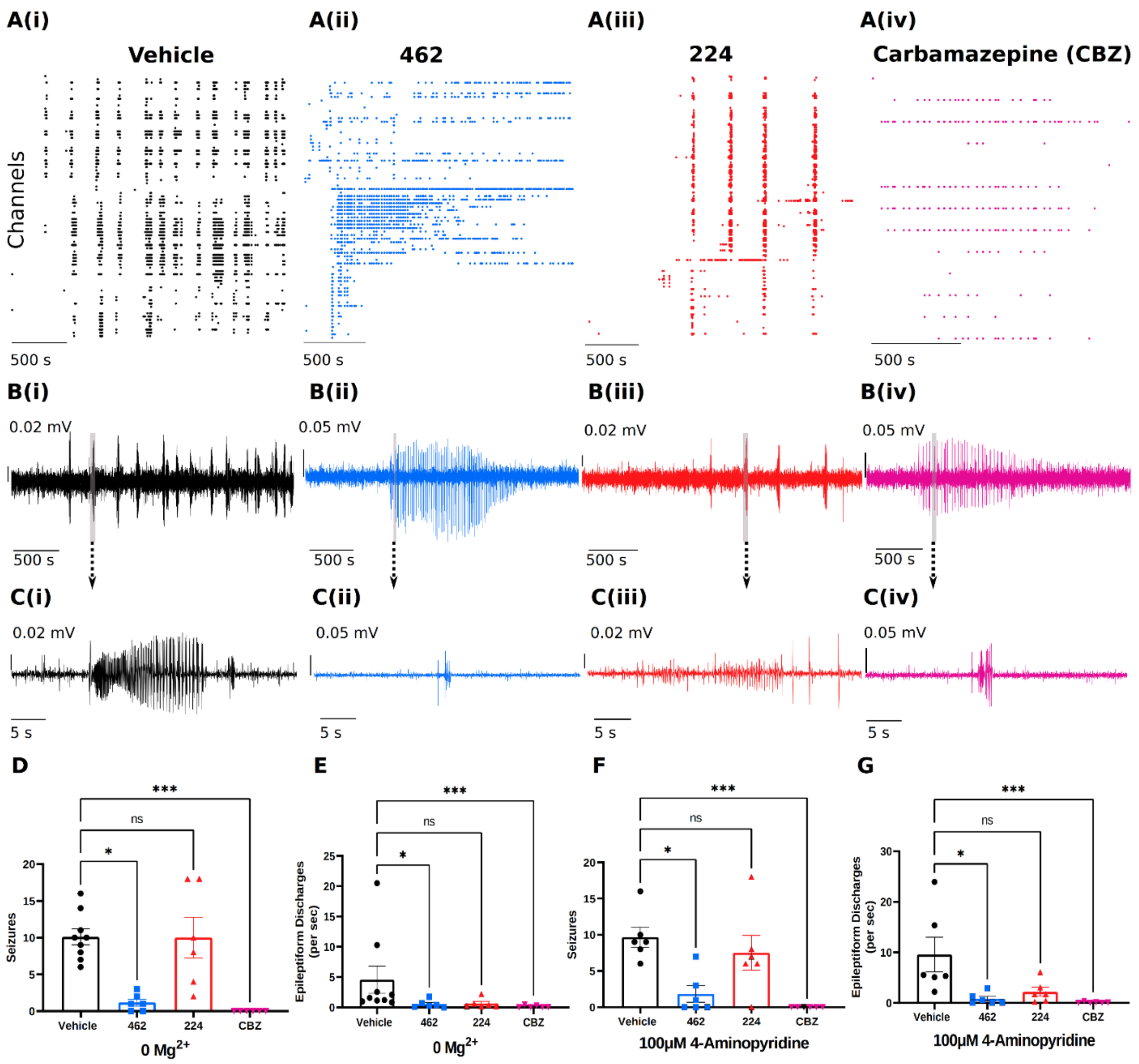


**Figure S5: Suppression of neocortical seizure-like events and epileptiform discharges in ex vivo seizure models by block of Na$_V$1.6 and Na$_V$1.2.** (A) Example raster plots from the 50-minute recording in 0 $Mg^{2+}$, 0 $Mg^{2+}$ + 150 nM XPC-5462, 0 $Mg^{2+}$ + 500 nM XPC- 7224, and 0 $Mg^{2+}$ + 100 µM CBZ. **(B)** Example traces of the local field potential activity from each condition. **(C)** Zoomed in view from the example traces, demonstrating the epileptiform activity observed. **(D)** 150 nM XPC-5462 and 100µM CBZ significantly reduced seizure-like events in the 0 $Mg^{2+}$ model of ictogenesis (Kruskal-Wallis test with Dunn's post-hoc test, N = 6-9). **(E)** 150 nM XPC-5462 and 100 µM CBZ significantly reduced total epileptiform discharges in the 0 $Mg^{2+}$ model of ictogenesis (Kruskal-Wallis test with Dunn's post-hoc test, N = 6-9). **(F)** 150 nM XPC-5462 and 100 µM CBZ significantly reduced seizure-like events in the 4-Aminopyridine model of ictogenesis (Kruskal-Wallis test with Dunn's post-hoc test, N = 6). **(G)** 150 nM XPC-5462 and 100 µM CBZ significantly reduced total epileptiform discharges in the 4-Aminopyridine model of ictogenesis (Kruskal-Wallis test with Dunn's post-hoc test, N = 6-9). (*P < 0.05, ***P < 0.001).
